# Chiral flows in the separating wall during cell division

**DOI:** 10.1101/2023.03.03.531016

**Authors:** Vijit Ganguly, Mainak Chatterjee, Anirban Sain

## Abstract

Material flow in the acto-myosin cortex of a cell, during cell division, has been found to be chiral in nature. Here we look for possible signature of such chirality during the growth of the intra-cellular membrane partition which physically divides the cell into two compartments. Many groups have recorded this partition formation phenomenon in C. elegans embryo, in real time, using fluorescent microscopy. We analyze some of these movies using PIV technique in order to search for signatures of chirality in the acto-myosin flow field on this partition. Further, we use standard hydrodynamic theory of active gell to predict possible chiral flow structures in the growing partition. While the flows in the growing annular shaped membrane partition is mainly radially inward, it can also develop non zero azimuthal velocity components due to chirality.

## I. INTRODUCTION

Many living organisms show chirality i.e., broken left-right (LR) symmetry, in their structure. For example, although in the exterior most animals appear symmetric, their spleen, liver, heart etc are placed on one side or the other of their midline. In lower eukaryotes, for example, shells show prominent chirality in their turns. In unicellular protozoa, cilia rotates in one direction and not both. Some of these are known to be manifestation of broken LR symmetry at the cellular level. In simpler multicellular invertibrates like Drosophila, c elegans etc the LR symmetry breaking of the cell is known to result in asymmetric embryogenesis and further differentiation of tissues. But it is not clear how this asymetry emerge from the chirality of the cell’s molecular building blocks, namely the DNA and the proteins which often form polymeric filament like structure. While all DNA molecules have helical conformation (hence chiral), exhibiting a particular handedness, some of the cytoskeletal proteins, like microtubule [1, 2] and actin [3], which are the main constituent of eukaryotic cell cortex are also made of helical arrangement of protofilaments. In some plants specific mutations have been shown to turn growth pattern from achiral to chiral, which is evidence of molecular level control of chirality [4]. Similarly, for cilia the direction of rotation can be directly mapped to the organization of actins. But for most other cases, specially in multicellular organisms, this link is unclear. One of the driving force behind large scale chirality could be chiral flow fields generated by cytoskeletal elements. For example, in some plants transport through vascular tissues are carried out by chiral flows driven by the helical motion of molecular motors along the walls [5]. In c elegans during the formation of cytokinetic actin ring chiral flows on the cortex towards the midcell have been detected [6]. Often both type of motions (right or left handed) can be detected in the same organism and they serve different purposes [5]. However one can also imagine local small scale chiral flows generating swirling motion which may not extend to macroscopic scales. Here we focus on one such possibility, namely, chiral motions in the separating wall during cell division.

A little background about cytoskeleton is perhaps needed here. Cytoskeleton is a thin mesh like structure made of actin filaments which are cross-linked by myosin motor proteins. It forms a layer beneath the cell membrane (cortex) providing mechanical stability to the cell shape. But these filaments undergo continuous poly/depoly-merization (turnover) and they can slide past each other as the crosslinks are transient. This makes the acto-myosin network self-regenerating and dynamical, yet maintaining actin alignment over longer length scales. Concerted mass movement in the cortex has been shown to cause change of cell shape, during cell division [6]. Considering the cortex to be a slow flowing gel, showing visco-elastic properties as well as filament ordering, has been an useful approach in deciphering the physics of cell deformation [7]. Further addition of inherent/active chiral torque to this theory has allowed an opportunity to study chiral motion in cell cortex. However the source of chiral torques at the molecular level leading up to macroscopic flows is not clear. In some cases, molecular motors have been found to serve as the link between the polar structure of the filaments and generation of macroscopic chiral flows. For example, Interactions between motors and elastic filaments (cilia) generate helical motion of the cilia which drive fluid flow [8] in the medium and helping locomotion in ciliary organisms. Actin polarity and cross linking molecules alpha-actinin have been linked to chiral rotation of the nucleus in certain cells [9]. Often cells appear to have inherent forces (torques in particular) which can cause rotations of both kind, i.e, clockwise and anti-clockwise. For example, a) with over expression of alpha-actinin-1 direction of nuclear rotations in cells were found to reverse in Ref [9]. b) large scale chiral motion could be induced in the Xenopus egg cell cortex by certain microtubule targeted drugs which caused preexisting f-actin fibers to re-organize and generate torsion forces [10]. This implies that emergence of a unique chirality in a class of organism may just depend on which torque is dominant.

In this paper we consider effect of active chiral torques in the actomyosin gel constituting a growing annular shaped cell-cell interface, with inner and outer radii *R*_0_ and *R*_1_, respectively (see Fig.1). We examine the emergent flow and alignment pattern of actin using a velocity field 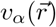 and nematic order parameter (OP) field defined as 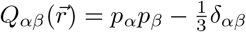, where *p*_*α*_ denotes polarity of the filaments. The actomyosin contractility driven interface closure, involving azimuthaly symmetric (i.e., *θ* independent) fields *Q*_*αβ*_(*r*) and *v*_*r*_(*r*) (along with *v*_*θ*_ = 0) was already discussed in our previous work [11]. It differed from the conventionally held view that the filament alignment in the actin ring is maintained by hydrodynamic flow [12]; instead we invoked recent experimental findings [13] that the observed rapid alignment is result of certain local, fast, directed polymerization processes. In our hydrodynamic theory this amounted to setting perfect alignment boundary condition at the inner edge of the closing interface. The resulting spatial actin alignment profile defined the ring thickness and accounted for enhanced active stress [11]. The present work focuses on effects arising from active body torques, e.g., emergence of nonzero *v*_*θ*_(*r*) field and corresponding changes in actin alignment.

**FIG. 1:**
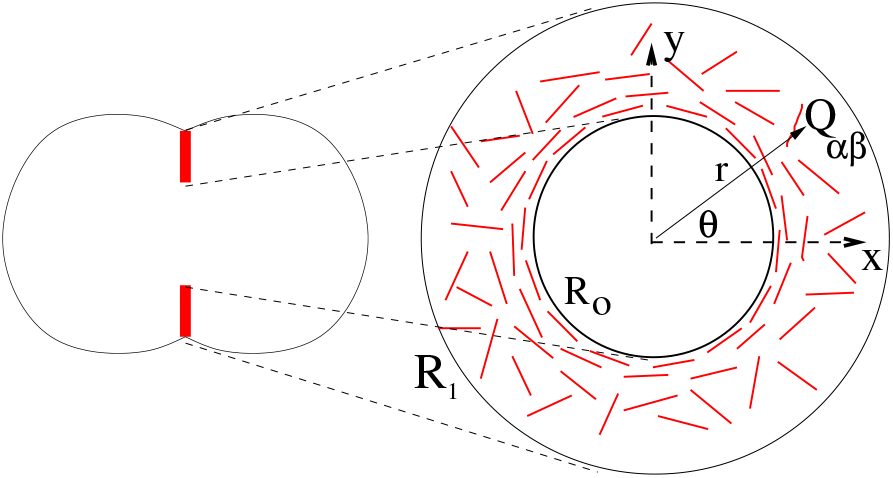
(a) The growing, annular shaped cell-cell interface (in red) during cytokinesis. (b) Cross-sectional view of the interface schematically showing organization of actin filaments (red lines) and quantitatively expressed in terms of the tensor order parameter field *Q*_*αβ*_(*r, θ*). The motion of the actomyosin gel on the annulus is expressed by the velocity field *v*(*r, θ*). The inner radius *R*_0_ shrinks with time as cytokinesis progresses, while the outer radius *R*_1_ remains fixed.

The paper is organized as follows. In section-1 we report PIV analysis of the velocity field obtained from movies of cytokinesis which captures closure of the interface in real time [14]. Section-2 briefly outlines the formalism which had been used before [11, 12] to describe the coupled dynamics of 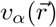 and 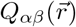 in the annular shaped growing interface. In section-3 we obtain the dynamical equations of the flow and the order parameter field, in the presence of active, chiral forces. In section-4 we present numerical solutions of these equations and discuss our results.

## II. SEARCH FOR CHIRAL MOTION AT THE GROWING INTERFACE

We performed PIV analysis of real time movies of cytokinesis, where the closure of the hole was captured from end-on view(see Fig.2). For the analysis reported here we used a movie from Ref[14] on c elegans. We sourced similar movies from Ref[15, 16], but they were too noisy for reliable PIV analysis. We analysed the velocity patterns at certain time instants indicated in the figures. The velocity magnitudes vary over more than an order of magnitudes across the interface, exhibiting high velocities near the ring and low velocities near the outer periphery. Due to this large spatial variation of speeds very small velocities were not observable because the high velocities set the scale bars. Further spurious contributions occur from the hole and the outside dark region. Therefore we masked the central hole as well as the outside while analysing the physically relevant interface. Further for quantitative analysis of the velocity statistics across different velocity scales we separately focused on a region near the ring (shown within the dashed box in Fig.2-b) and the region away from the ring (outside the dashed circle shown in Fig.2-c. Note that the hole is not centrally located in the interface and is also not fully circular. So after obtaining the absolute velocity vectors through PIV in the cartesian frame we decomposed them into radial and azimuthal components *v*_*r*_ and *v*_*θ*_ after assuming the center of the approximately circular hole as the origin of the polar coordinates. The resulting histograms (unnonrmalised probability distributions) of *v*_*r*_ and *v*_*θ*_ are shown in Fig.3. The velocity showed high inhomogeneity across the interface, differing by an order of magnitude between the zones near the ring and zones away from it. We checked this by masking out the circular high intensity zone in Fig.2-c. For computing velocity histograms from the boxed zone in Fig.2-c masking out the central hole was not effective because the hole was not exactly circular in shape. However after collecting velocity data for histogram (Fig.3a,b) we ascertained that the dominant high peak near the zero was due to the spurious low velocity signal (noise) from the central hole. The insets in Fig.3 are drawn after removing this noise. Fig.3-a shows dominant radial inward flow, while the same exercise revealed weak signature of nonzero *v*_*θ*_ near the ring (the region in the box in Fig.2). Velocity vectors in the region away from the ring is shown in Fig.2.

**FIG. 2:**
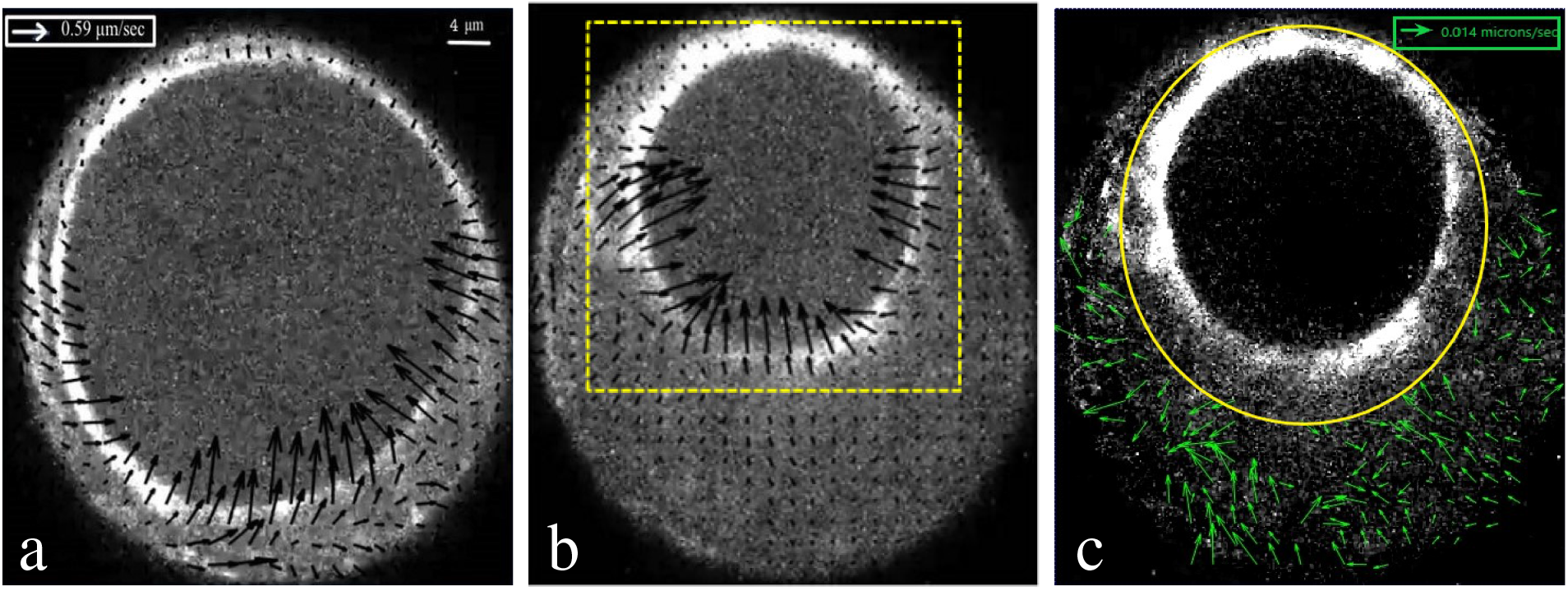
Velocity field (arrows) on the interface during ring closure, obtained from PIV analysis, (a) after 70 sec while (b) and (c) both after 105 sec after initiation of closure. The velocity near the ring (∼ 0.5*μ* m/s) is prominent in (b) and is more than an order of magnitude larger than that away from the ring which is shown in (c) by masking the region around the bright ring. The dark region (the hole) at the middle of the (a) and (b) are masked during PIV to avoid spurious velocity vectors. While plotting velocity distributions only the boxed region from (b) is used. Scale bars for distance are same across (a),(b),(c), while the velocity bars are same in (a),(b) and differs for (c).

**FIG. 3:**
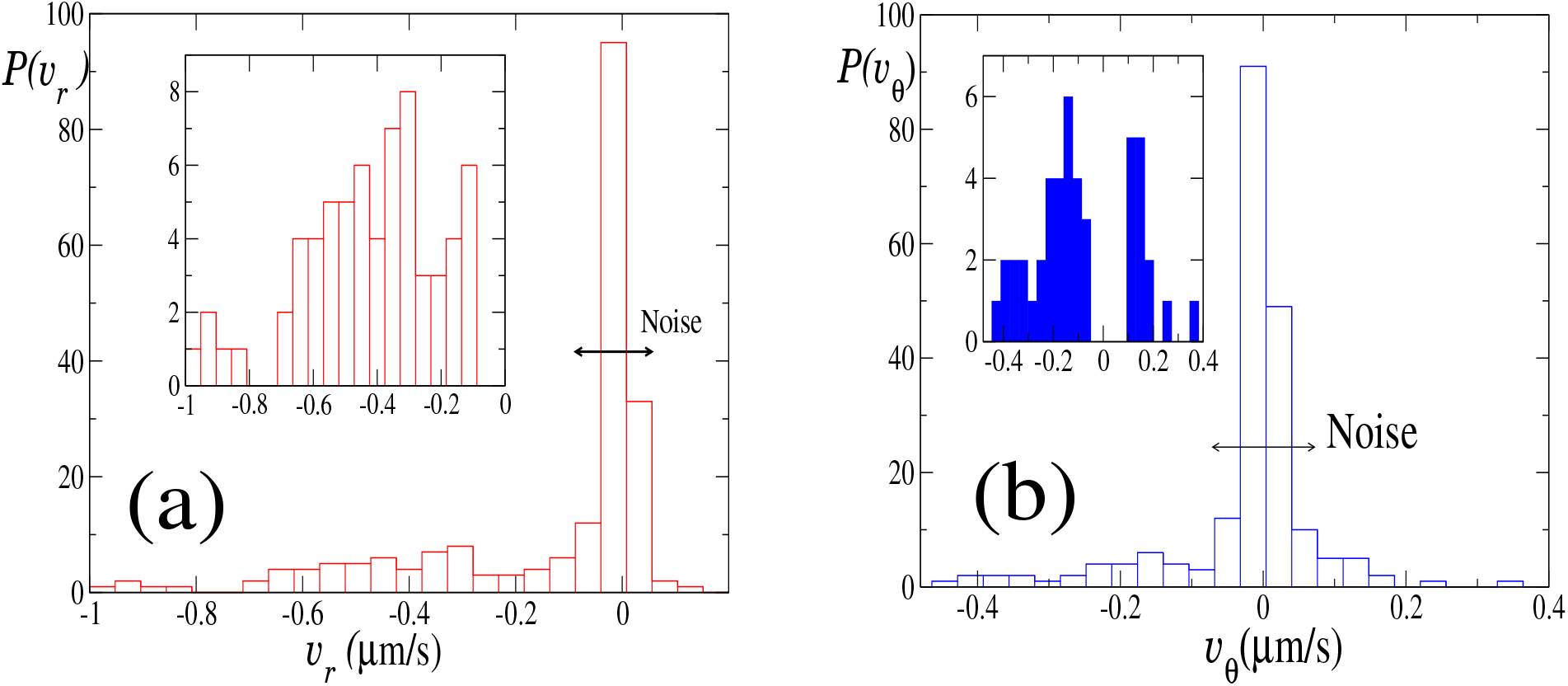
Probability distributions (unnormalized) of (a) radial velocity component *v*_*r*_ and (b) azimuthal velocity component *v*_*θ*_, obtained at *t* = 105 sec (Fig.2) using PIV analysis. PIV is done for the box shown in Fig.2. The high peaks near zero (marked by the horizontal double headed arrows) are identified with noise from the zones designated as ‘a’ and ‘b’ in Fig.4). The shallow peaks are identified with the bright zone in the box around the ring and highlighted in the corresponding insets. For *v*_*r*_ the distribution is highly skewed towards negative *v*_*r*_ implying the flow is mainly radially inward, but for *v*_*θ*_ it is relatively less skewed towards negative *v*_*θ*_. The histograms in the insets are replots of the respective distributions in the main figures after filtering out the noise near zero.

**FIG. 4:**
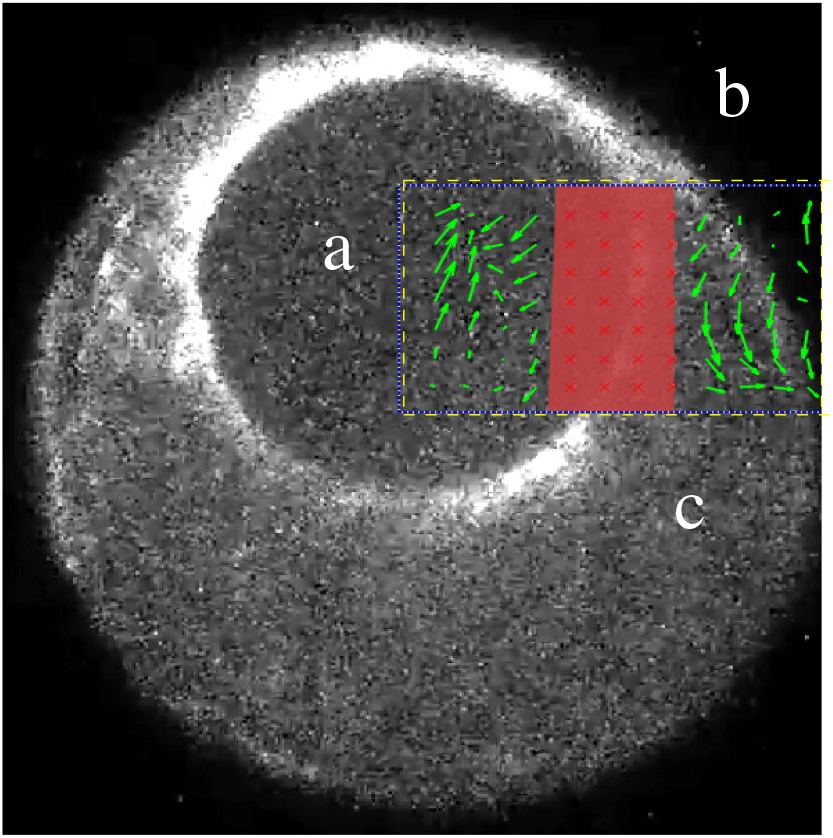
To asses the spurious velocity scales in the regions ‘a’ and ‘b’ we performed PIV analysis within the yellow box after masking the bright zone near the ring (pink box). It showed that the velocities in zones ‘a’ and ‘c’ are of order 0.04*μm*, while in ‘b’ they are even smaller.

## III. FORMALISM

For the non-chiral active forces, the details of the velocity and OP calculations pertaining to the growing interface are already available in Ref[11] and the Supplementary information there in. Here we will define the various terms involved in the dynamical equations [12], for completeness sake, and mainly discuss the changes due to active torques. We start with an isotropic arrangement of nematics, in the 2D x-y plane of the interface, in the absence of flow or anchoring boundary conditions. The corresponding order parameter matrix is 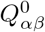. Due to tracelessness and isotropy in x-y plane, it is diagonal with 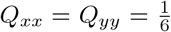 and 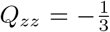. In the presence of flow and boundary condition at the inner edge (*R*_0_) the OP changes to 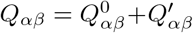. Again due to tracelessness and the fact *p*_*z*_ = 0, the only nonzero components of 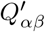 are 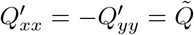 and 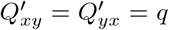. Note that these are average values of the *Q*_*αβ*_ components.

The constitutive equations of the stress and order parameter evolution in active gel are [12]

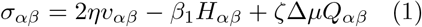

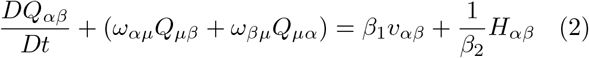

Here in Eq.2, 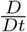 is the material derivative and the 2nd term on the left hand side is the co-rotational derivative involving vorticity 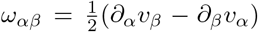. In the steady state the whole left hand side is set to zero. Further, *σ*_*αβ*_ consists of an elastic part 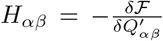 based on the nematic Landau–de Gennes free energy 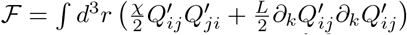. This form favours an isotropic arrangement of the nematic with a correlation length 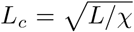. Due to strong boundary alignment of the nematics at the inner peripheri (at *r* = *R*_*0*_) the alignment turns out to be strong near *r* = *R*_*0*_ (the so called actin ring) and decays exponentially away from it with the length scale *L*_*c*_, which can be interpreted as the effective width of the actin ring [11]. The stress also has a part proportional to the viscous strain rate 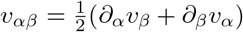 and an anisotropic active component *ζ*Δ*μQ*_*αβ*_, where Δ*μ* is the chemical potential difference generated due to ATP hydrolysis. Here *η* is the fluid viscosity and *β*_*1*_ is the flow coupling that affects 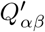. The coefficients *β*_*1*_ and *β*_*2*_ are Onsager coefficients [17, 18] that relates the fluxes *σ*_*αβ*_ and 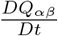, to the conjugate forces *H*_*αβ*_ and *v*_*αβ*_ by linear relations [17]. Further, following Ref[12] we assume the interface thickness (*e*) to be small and constant, and define a 2D effective tension *t*_*ij*_ = *e*(*σ*_*ij*_ − *δ*_*ij*_*σ*_*zz*_), where we have used pressure *P* = *σ*_*zz*_, assuming the net normal stress on the interface to be zero. Using this tension tensor we work with a 2D hydrodynamic theory where the force balance equation is 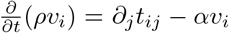. Here *α* is the cytoplasmic friction tangent to the flat, annular shaped interface (see Fig.1) with a shrinking inner radius *R*_*0*_(*t*) and a fixed outer radius *R*_*1*_. respectively. Using 2D polar co-ordinates, and in the highly viscous regime, the force balance equations are 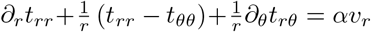, and 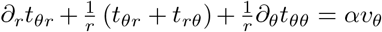. In the 2D polar coordinates the 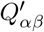 matrix changes to 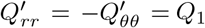 and 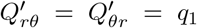, i.e., retains the symmetric traceless structure [11]. Here we consider the velocity and order parameter fields to be functions of *r* only and independent of *θ* (azimuthal symmetry). Therefore all derivatives with respect to *θ* are dropped; however we do not assume *v*_*θ*_ = 0. Without any chiral term in the stress field it turns out [11] that the dynamics of *v*_*θ*_ is given by (ignoring friction)

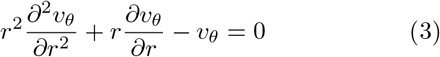

Eq.3 has a general solution, 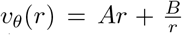. With boundary conditions *v*_*θ*_(*r* = *R*_*1*_) = 0 at the outer periphery and zero shear stress at the free inner boundary *σ*_*rθ*_(*r* = *R*_*0*_) = 0, one can show that, *v*_*θ*_ = 0. In the next section we will consider the effects of nonzero chiral terms in the constitutive equations.

## IV. THEORY WITH ACTIVE TORQUE

Our PIV analysis of the cortical flows on the dividing interface shows weak non-zero azimuthal component i.e., *v*_*θ*_ ≠ 0. This could be due to the asymmetry in positioning of the hole during its closure, i.e., the hole may not be at the center of the dividing interface, which has a nearly circular outer periphery. This asymmetry can drive non-radial flows as was shown in Ref.[14]. In fact such asymmetric closure is more common than symmetric radial closure when only *v*_*r*_ is nonzero. But intrinsic chirality at the molecular level can also generate nonzero *v*_*θ*_ at a larger scale. Chiral flows have been observed on the cortex of c elegans during the formation of the ring at the equator. Flows originating at the poles of the ellipsoidal shaped cell and flowing towards the equator develop azimuthal components [19] (parallel to the equator). As a result flows from the two opposite poles can enter the equator generating a twist at the equator. Further Ref[6] has detected and analysed chiral flows on the cortex. Given that the growing cell-cell interface is also made of cytoskeletal acto-myosin network it is likely to share the same properties with active cell cortex, including its chirality. Therefore it is natural to ask about the consequence of chirality, in the flows on the growing interface, which we now discuss in details.

Several papers [12, 18, 20, 21] have shown how existence of chirality can modify both the symmetric and anti-symmetric part of the stress tensor. In the absence of external torques and body torques local rotational equilibrium dictates that stress tensor be symmetric. But inclusion of body torques can render the antisymmetric part of the stress tensor non-zero in addition to modifying the symmetric part as well. Ref [18] has shown how microscopic origin of the body torque can lead to different terms to the hydrodynamic stress tensor.

The chiral effects on the hydrodynamic flow and order parameter field which we report here is due to addition of a chiral term to the symmetric part of the stress tensor, similar to that used in Ref[22], albeit in a different context. We will first summarize our derivation of this additional chiral term in the stress. We will mainly follow the formalism of Ref[18], with minor change of notations. For example, we express angular momentum vector *l*_*α*_ by its dual representation *l*_*αβ*_ = *ϵ*_*αβγ*_*l*_*γ*_ and similarly, the angular velocity tensor as Ω_*αβ*_ = *ϵ*_*αβγ*_Ω_*γ*_ where Ω_*γ*_ is the angular velocity vector. To distinguish the active terms from their passive counterparts we use the superscript ′*A*′.

We consider an incompressible fluid made up of identical rigid rods (uniaxial nematics) 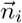 with their center of masses (CM) at 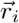, with respect to the lab frame and where *i* denotes the *i*-th rod. Each rod rotates about its own CM with angular velocity Ω_*i*_. Thus spin angular momentum is *l*_*i*_ = *I*Ω_*i*_, where *I* is the moment of inertia tensor of the rod. The hydrodynamic variables used are coarse grained average of these microscopic variables. The total angular momentum density of the fluid is 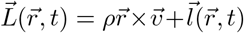, sum of orbital and the intrinsic spin part. Here 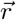 is the hydrodynamic position of a small volume element *dV* which consists of many identical rods at positions 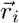.

The dynamics of the intrinsic spin is governed by passive and active torques, denoted as *T*_*αβ*_ and 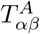, respectively.

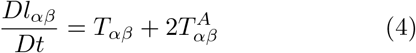

Here, 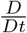 is convective derivative and the factor of two in the active term is a matter of convention. We will not consider any surface contribution to the passive torque as it gives rise to higher order terms in derivatives [18]. Thus *T*_*αβ*_ is the body torque while the active torque 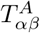 includes both surface and body torques (see below).

Using conservation of total momentum (*ρv*_*α*_) and total angular momentum 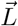, in the absence of any external forces and torques, one can show that the passive body torque 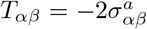 [18, 20], where ‘a’ denotes the antisymmetric part of the stress. Since *T*_*αβ*_ and *l*_*αβ*_ both are anti-symmetric, Eq.4 dictates that 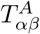 must also be anti-symmetric. Note that *σ*_*αβ*_ is purely passive, which verifies the standard result that at equilibrium, absence of unbalanced torques (*T*_*αβ*_) inside a body dictates that the antisymmetric part of the stress tensor must be zero and therefore the net stress *σ*_*αβ*_ is completely symmetric. We assume the active body torque 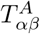 to have both surface and volume contributions. The surface part is represented by a divergence term, 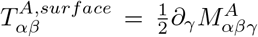 and the volume term by 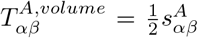. Thus Eq.4 reduces to

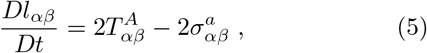

where,

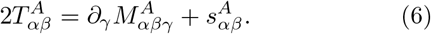

In the steady state, 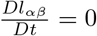 we have

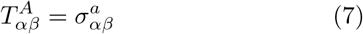

The passive part of stress 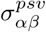 has additional contributions over its non-chiral counterpart. Existence of nonzero relative spin Ω_*αμ*_ − *ω*_*αμ*_ brings in rotational viscosity terms which are separated into symmetric and antisymmetric parts (indicated by superscripts ‘s’ and ‘a’, respectively :

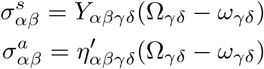

The dissipation coefficients can be functions of nematic order, generating coupling between the relative spin and the local nematic order [18]. For example,

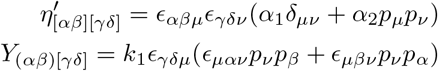

Following the notations in Ref[18], here round ‘()’ and square ‘[]’ brackets indicate symmetric and antisymmetric pair of indices, respectively, while *α*_*1*_, *α*_*2*_, *k*_*1*_ are constants. Here we have retained terms that are of lowest order, individually, within symmetric and antisymmetric terms. For example, for *η*′ we retain only the isotropic term (involving *α*_*1*_). The second term involving *α*_*2*_ can generate an anti-symmetric term quadratic in *p*, similar to the ones in *Y*_(*αβ*)[*γδ*]_ but we do not include this here for simplicity.

After simplifying and combining the dissipative and reactive terms, the net stress reads [18]

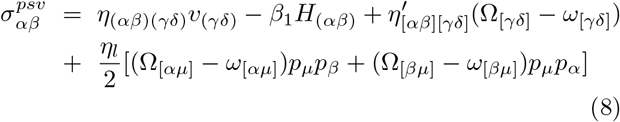

For clarity, we have indicated the symmetric and antisymmetric pair of indices, which we will further omit. Note, while *v*_*γδ*_ is symmetric in *γδ*, the relative angular velocity (Ω_*γδ*_ − *ω*_*γδ*_) is anti-symmetric. Therefore, the respective viscosities *η*_*αβγδ*_ and 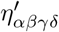 are symmetric and anti-symmetric in *γδ*, respectively. The *η*_*l*_ term is symmetric in *α, β*.

For simplicity we consider both the viscosities to be isotropic. Thus the standard shear viscosity term becomes *ηv*_*αβ*_ and the using 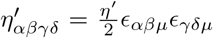 the rotational viscosity term reduces to 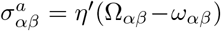. We now substitute this anti-symmetric part of passive stress into eq(7) which yields,

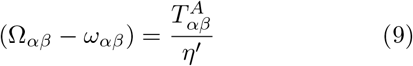

Including the active stress 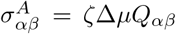, as in Eq.1 the force balance equation is

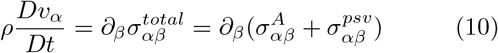

Using Eq.9 for the relative spin terms in the passive stress (Eq.8) the right hand side of the above equation reduces to

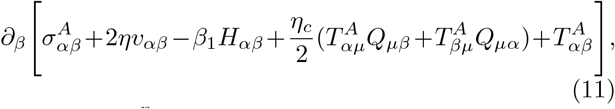

where 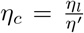. The rest of the terms in the net stress above can be attributed to chairality i.e., existence of local intrinsic body torque which gives rise to a relative spin to the directors over and above the local fluid vorticity. Thus

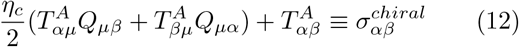

In a similar fashion adding chirality to the director equation (2) we get,

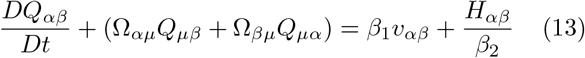

which upon using Eq.(9) reduces to,

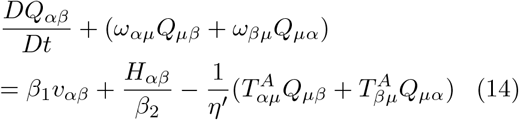

Note that *Q*_*αβ*_ being symmetric in *αβ*, the active terms have been constructed to preserve this symmetry. Further, we need to determine the active term 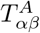 asymmetric in *αβ*. As we already saw in eq(6), 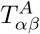 consists of a bulk term 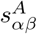 and an antisymmetric surface term 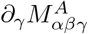 Following Ref[21]),

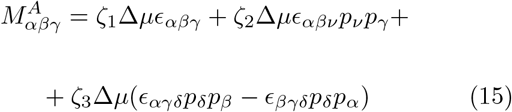

Since the nematics in our system mainly lies in *xy* plane with a negligible *p*_*z*_ component, *γ* = *z*. Further *α* ≠ *β* and they are either (*x, y*) or (*y, x*). Under one constant approximation (*ζ*_*1*_ = *ζ*_*2*_ = *ζ*_*3*_ = *ζ*) we get 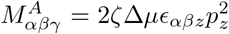. Thus 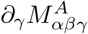 vanishes under the limit *p*_*z*_ → 0. The remaining active bulk stress 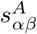 is also antisymmetric and is expected to be function of *p*_*γ*_ as well.

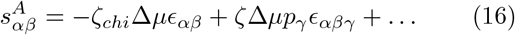

As we discussed earlier, (*α, β*) being (*x, y*) or (*y, x*), and *p*_*γ*_ = *p*_*z*_ being zero, the 2nd term vanishes, and we take the 1st term as the simplest choice for 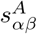, and thus, 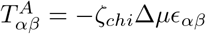. Here the minus sign of the isotropic term is a mere convention. With this choice of spatially uniform 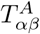 the antisymmetric part of the total stress (the expression within square brackets in Eq.11) does not contribute to the force balance equation (as spatial derivative of 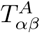 is zero). However, 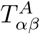 contributes via its coupling to the nematic order parameter (i.e., the *η*_*c*_ term in Eq.11) which is symmetric in (*αβ*). Thus the net effect of chirality on the constitutive equation is an additional symmetric term in the stress. Substituting the expression for 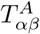 the net stress and the director equations read,

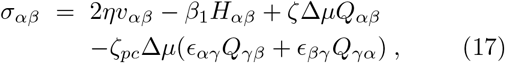

where 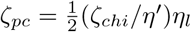, and

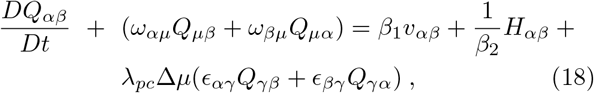

where *λ*_*pc*_ = *ζ*_*chi*_*/η*′.

Eq.18 and the force balance equation, resulting from Eq.17, are coupled non-linear equations of the velocity *v*_*α*_ and the order parameter field *Q*_*α,β*_. We will solve these numerically (using Mathematica) with appropriate boundary conditions. But first we will look at an over-simplified limit where we will ignore the effect of velocity on the OP. We set *β*_*1*_ = 0 and the symmetric coupling between *ω*_*αβ*_ and *Q*_*αβ*_. This allows a analytic solution of the *Q*_*α,β*_ field in the steady state 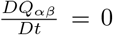, which we substitute into the force balance equation to obtain the *v*_*α*_ field. This gives us a gross feel about the effect of the active chiral term (*λ*_*pc*_) on the OP field. We impose a phenomenological boundary condition that the nematic field at the inner boundary is fully aligned with the boundary. It turns out that this boundary condition dominates the solution of the nematic field even when the velocity dependent terms are present. Choice of this boundary condition is motivated by the recent discovery [13] that there are fast, local, directed polymerization processes (much faster compared to the aligning effect of the velocity field) which generates and maintains the actin alignment in the actin ring at inner boundary. Therefore on the slow time scale of these hydrodynamic equations for *v*_*α*_ and *Q*_*αβ*_ the nematics at the boundary is always aligned [11].

After ignoring all the velocity dependent terms in Eq.18 and assuming steady state 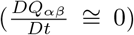, we get *H*_*αβ*_ in terms of the chiral term.

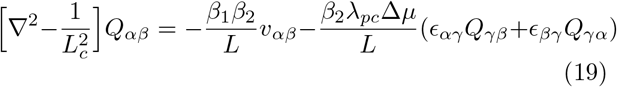

where the expression for − *H*_*αβ*_, evaluated from the Frank free energy, appears on the left hand of the equation. The diagonal and the off diagonal terms of this equation yield two independent equations for 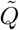 and *q*. We recall that 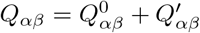 and 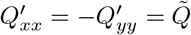 and 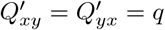

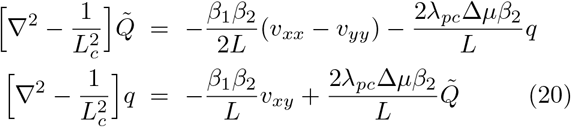

While deriving these equations the 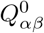 terms do not survive due to their constant values and symmetric form. **check**. These two equations show how chirality affects different components of order parameter. In the polar form these equations changes to the following form, with *Q*_*1*_, *q*_*1*_ being the polar counterparts of 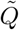, *q* [11].

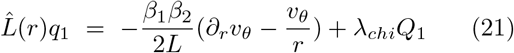

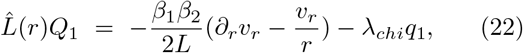

where the operator 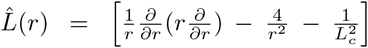 and *λ*_*chi*_ = 2*λ*_*pc*_Δ*μβ*_*2*_*/L*. The force balance equation 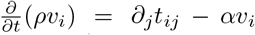 in the low Reynold’s number regime (where the inertial term is ignored) reduces to, ∂_*j*_*t*_*ij*_ = *αv*_*i*_. In the polar coordinate frame it leads to two independent equations. The one involving radial velocity *v*_*r*_(*r*) reads,

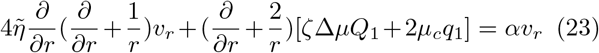

The azimuthal component of force balance equation involving *v*_*θ*_(*r*) reads,

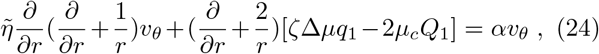

where 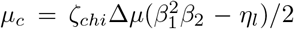 and 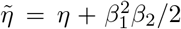 is the renormalized viscosity. Note that the sign of *μ*_*c*_ does not get fixed with a chosen sign of *ζ*_*chi*_ i.e., a chosen handedness. The relative magnitudes of flow coupling parameter *β*_*1*_, the elastic relaxation time-scale *β*_*2*_ of the nematic field and viscous coupling *η*_*l*_ between the relative local rotation and the nematic field, together decide the sign of *μ*_*c*_ which determine the resulting azimuthal flow *v*_*θ*_.

Eq.22-24 are our main set of equations for chiral dynamics and we solve them simultaneously, numerically with appropriate boundary conditions. At the outer boundary *r* = *R*_*1*_, we set *q*_*1*_ = 0, *Q*_*1*_ = 0, *v*_*r*_ = 0, *v*_*θ*_ = 0. The inner boundary, at *r* = *R*_*0*_, is stress free and thus *σ*_*rr*_ = *σ*_*rθ*_ = 0. The actin (nematics) is assumed to be completely aligned with the inner peripheri and thus *Q*_*1*_ = − 1*/*2 and *q*_*1*_ = 0.

Plots of *Q*_*1*_, *q*_*1*_, *v*_*r*_(*r*), *v*_*θ*_(*r*) are shown in Fig.5. Some interesting features emerge from these solutions. First, in the absence of external torques or explicit velocity boundary conditions (for example, azimuthal velocity *v*_*θ*_(*R*_*1*_) ≠ 0 at the boundary), nonzero *q*_*1*_ and *v*_*θ*_ fields can still arise due to intrinsic chiral active stresses. We checked this by solving Eq.22-24 numerically after setting *λ*_*c*_, *μ*_*c*_ = 0. On the other hand, for *λ*_*c*_, *μ*_*c*_ ≠ 0 the resulting magnitude and direction of *v*_*θ*_ depend on the magnitude and sign of *λ*_*c*_ and *μ*_*c*_. Interestingly, for the same sign for *ζ*_*chi*_, the sign of *v*_*θ*_ can reverse depending on the sign of 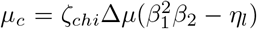 which can be seen by comparing Fig.5-d and Fig.7-d, where the signs of *μ*_*c*_ are positive and negative, respectively. However the *Q*_*1*_, *q*_*1*_ and *v*_*r*_(*r*) fields do not qualitatively change, only their magnitudes are modified (compare a,b,c in Fig.5 and Fig.7. This implies that the molecular level chirality *ζ*_*chi*_ does not uniquely decide the direction of the azimuthal flow *v*_*θ*_ but it depends on the relative strengths of the couplings. This opens up the possibility that depending on inhomogeneity of coupling strengths different rotation directions may be visible in the same system.

**FIG. 5:**
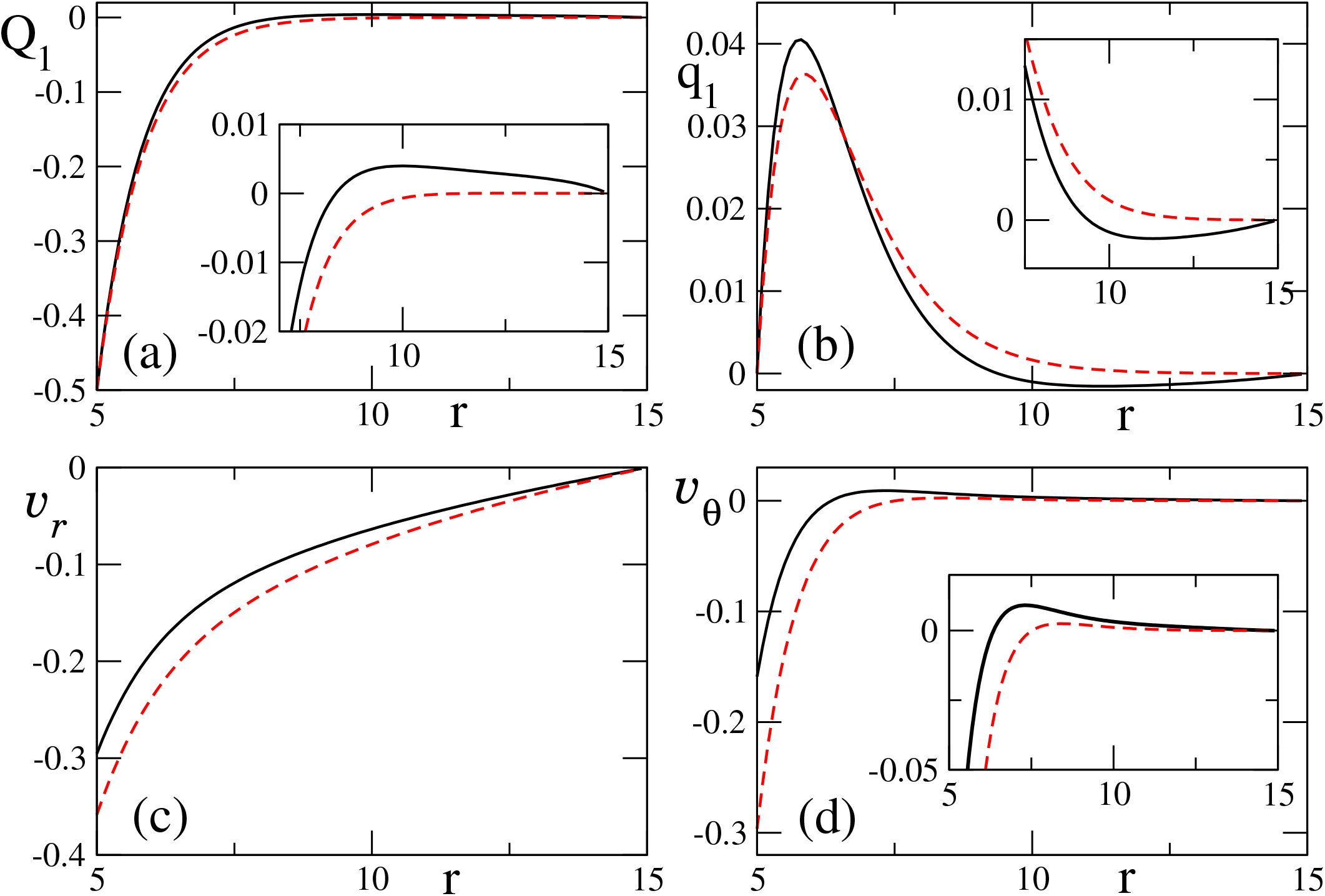
Spatial (*r*) variations of the order parameters a) *Q*_1_, b) *q*_1_ and the velocities c) *v*_*r*_, d) *v*_*θ*_ are shown. In all the figures the dashed lines are for *β*_1_ = 0, i.e., when the order parameters *Q*_1_, *q*_1_ are not not influenced by the velocity field (Eq.22. The solid lines are for *β*_1_ ≠ 0 case and in sub-figures a,b and d, they reveal a nonmonotonic approach to zero at the outer peripheri *R*_1_. We have provided insets in these sub-figures to highlight this feature. This nonmonotonicity, interestingly results in the reversal of direction of the azimuthal flow *v*_*θ*_ at some finite radius. The parameters used are *R*_0_ = 5*μm, R*_1_ = 15*μm, L*_*c*_ = 1*μm, L* = 1.0*μm, e* = 1*μm, η* = *β*_2_ = 1.0, *β*_1_ = 0.5, *ζ*Δ*μ* = 1.0 and the chiral parameters *λ*_*chi*_ = 0.5, *ζ*_*pc*_ = 0.5, *μ*_*c*_ = −0.37.

In Fig.6 we show the velocity field vectors corresponding to the solutions shown in Fig.5. Note the chiral *v*_*θ*_ components which is more prominent near the inner boundary. However from the information of *Q*_*1*_ and *q*_*1*_, we get 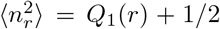 (and consequently, 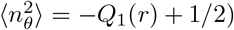, and ⟨*n*_*r*_*n*_*θ*_⟩ = *q*_*1*_, we cannot compute the corresponding averages ⟨*n*_*r*_⟩ and ⟨*n*_*θ*_⟩ because the underlying probability distribution is not known. From the boundary conditions, we can infer that, at the inner boundary (*r* = *R*_*0*_), ⟨*n*_*θ*_⟩ = 1 and ⟨*n*_*r*_⟩ = 0. Similarly, at the outer boundary (*r* = *R*_*1*_) we must have both ⟨*n*_*θ*_⟩ and ⟨*n*_*r*_⟩ = 0. The nematic orientations shown in schematic Fig.1 is consistent with these boundary conditions, except that the non-zero *q*_*1*_(*r*) is not reflected in this schematic.

**FIG. 6:**
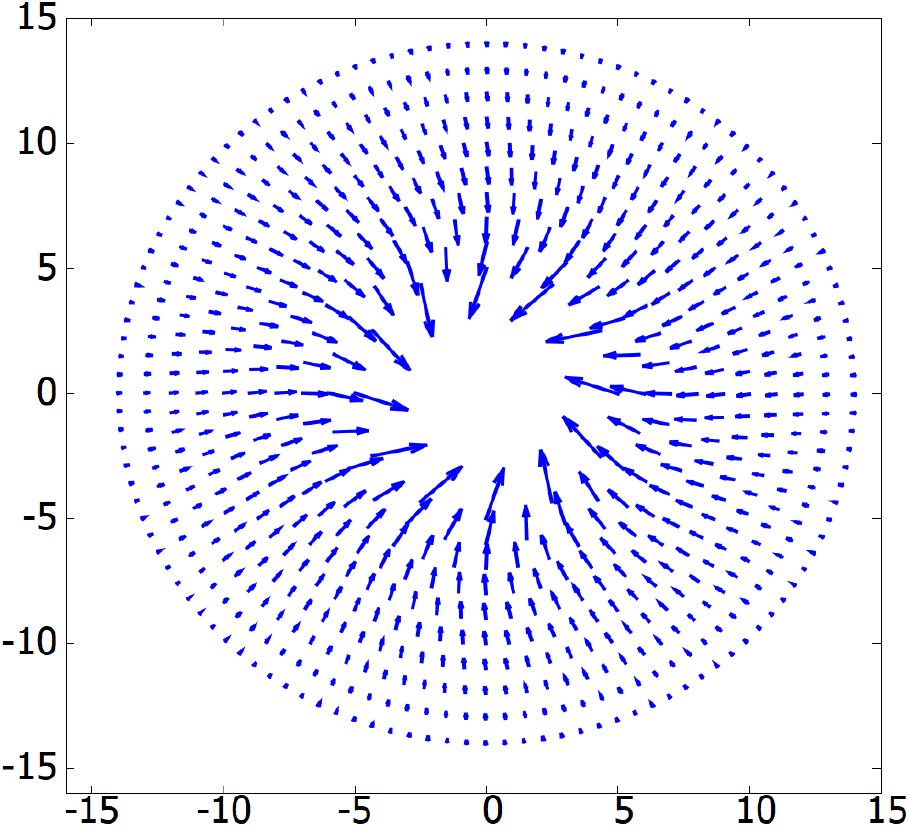
Velocity field obtained from (*v*_*r*_, *v*_*θ*_) of Fig.5. Note the weak *v*_*θ*_ component prominent near the inner boundary of the interface.

Second, we note that all the plots, except that of *v*_*r*_ show non-monotonicity with respect to radial distance *r*. Since *Q*_*1*_, *q*_*1*_, *v*_*r*_, *v*_*θ*_ are all dependent on each other via Eq.22-24 their nonmonotonicity will be inter related. So far *q*_*1*_ is concerned, it is set to zero at the inner boundary assuming complete tangential alignment of the nematics there. Similarly, at the outer boundary *R*_*1*_ it is set to zero due to the assumption of isotropic orientation there. Therefore any nonzero solution for *q*_*1*_ will be non-monotonic. However, the solution in Fig.5-b shows a weak non-monotonicity as it approaches zero at large *r* which is an emergent effect and the reason for this is not clear. Interestingly, for *v*_*θ*_ this non-monotonicity amounts to change of direction with increasing *r*, from clockwise to anti-clockwise or the reverse, depending on the sign of *μ*_*c*_.

Third, note that coupling of *Q*_*1*_, *q*_*1*_ to the velocity field via *β*_*1*_ increases the effective viscosity (as noted below Eq.23,24) which damps the velocity field (*v*_*r*_, *v*_*θ*_). Also, for these plots we ignored cytoplasmic friction (the *α* terms on the right hand sides of Eq.23,24) which are known to further damp the velocity field [11].

We also computed *σ*_*θθ*_ as a function of *r* (not shown here). *σ*_*θθ*_ is the main driving force for constriction. It is always positive but shows non-monotonic behaviour qualitatively similar to that in the absence chirality as seen in Ref [11]. The fact that *σ*_*θθ*_ always remain positive is reflection of contractility, which is highest at the ring. However the reason for its non-monotonicity is not clear.

## V. CONCLUSION

The formal part of this work mainly involved finding how active chiral torque 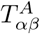 modify the net stress field. Eq.11 shows that modifications arise both in the symmetric and antisymmetric part of the stress. However, due to our simplest choice 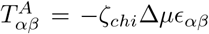 which a spatially uniform, the asymmetric part did not contribute to the dynamics, since the force balance equation depends only on derivative of *σ*_*αβ*_. On the other hand, due to Onsager symmetry relations 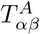 couple symmetrically with the OP field (see Eq.11) which is spatially inhomogeneous with nonzero spatial derivative and hence contribute to the dynamics (Eq.10). Note that the nonzero *v*_*θ*_ that emerged here is explicitly due to the active chiral stress tuned by *ζ*_*chi*_. However the direction of *v*_*θ*_ is controlled by the sign of [*ζ*Δ*μq*_*1*_ − 2*μ*_*c*_*Q*_*1*_] in Eq.24 which depends on the relative strengths of activity *ζ*Δ*μ*, the flow coupling parameter *β*_*1*_, rotational viscosity *β*_*2*_ and viscous coupling *η*_*l*_ between the relative local rotation and the nematic field. Note that for the non-chiral case (*ζ*_*chi*_ = 0) both *q*_*1*_ and *μ*_*c*_ are individually zero, leading to *v*_*θ*_ = 0. But when *ζ*_*chi*_ ≠ 0 and yet *μ*_*c*_ = 0 we can have nonzero *v*_*θ*_ due to nonzero *q*_*1*_. In our calculation *Q*_*1*_ was an order of magnitude larger than *q*_*1*_ and therefore the sign of [*ζ*Δ*μq*_*1*_ − 2*μ*_*c*_*Q*_*1*_] was mainly controlled by *μ*_*c*_ and we examined reversal of flow direction in response to changing the sign of *μ*_*c*_. We note that Ref[22] did also encounter this feature while trying to model cell orientations and flow in a parallel channel, and there sign of *μ*_*c*_ could be fixed by comparing with the preferred angle of the cells in the channel. However, in our present case, for flows in the cell-cell interface, which is at much smaller scale compared to the tissue flow, no such clear experimental results are available yet on the direction of *v*_*θ*_. Therefore our PIV analysis was limited to only one movie and it showed weak signature of *v*_*θ*_. We speculate that if [*ζ*Δ*μq*_*1*_ − 2*μ*_*c*_*Q*_*1*_] is close to zero then small temporal fluctuation (noise) of the coupling parameters, discussed above, may lead to different rotation directions at different times, which could be a reason for not detecting unidirectional motion. The weak reversal of *v*_*θ*_ along the radial direction (Figs.5-d and 7-d) is also interesting and we did not find any precedence of such motion in the literature.

**FIG. 7:**
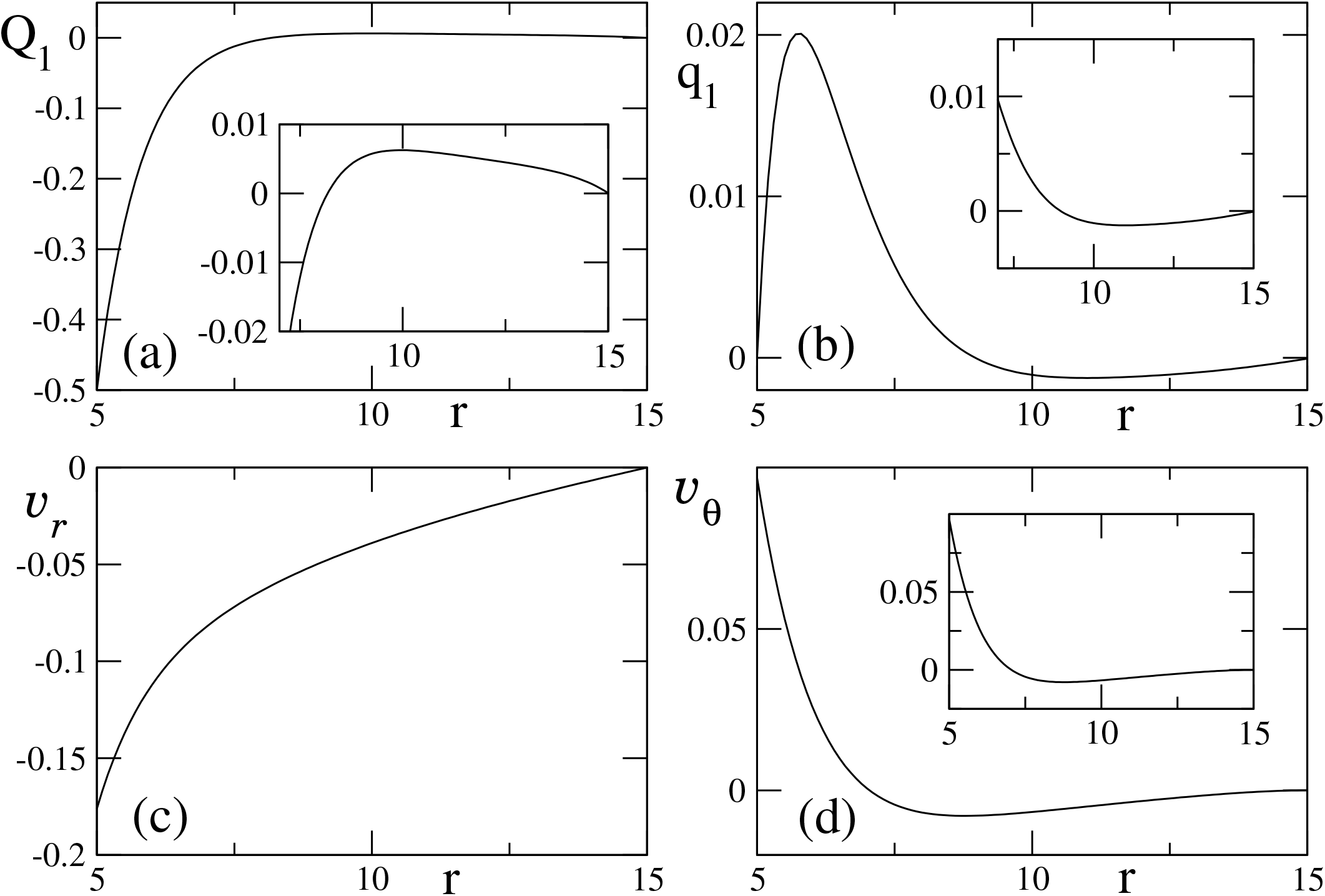
Change of sign in *μ*_*c*_ reverses the direction of the azimuthal flow *v*_*θ*_(*r*) compared to that in Fig.5. This is shown in (d). However the spatial variations of *Q*_1_, *q*_1_ and *v*_*r*_ (shown in a,b,c and the accompanying insets) do not change qualitatively; only the magnitudes of *q*_1_ and *v*_*r*_ are reduced. The parameters are same as in Fig.5 except now *μ*_*c*_ = 0.22 and *β*_1_ = 1.2

